# EFFECT OF A BIORESONANCE DEVICE ON VIABILITY AND METABOLIC ACTIVITY IN HUMAN UMBILICAL VENOUS ENDOTHELIAL CELLS

**DOI:** 10.64898/2026.01.30.702890

**Authors:** Marco Cosentino, Emanuela Rasini, Marco Ferrari, Alessandra Luini, Massimiliano Legnaro, Franca Marino

**Affiliations:** Center for Research in Medical Pharmacology, University of Insubria, Varese (I)

**Keywords:** bioresonance, human umbilical venous endothelial cells, viability, metabolic activity, QDome Mini Personal

## Abstract

The present study was aimed at evaluating the effects of the bioresonance (BR)-based device QDOME MINI PERSONAL, produced by the society QMED SWISS SA (Lugano, CH), on the viability and mitochondrial metabolic activity of cultured human umbilical venous endothelial cells (HUVEC).

To this end, HUVEC were cultured under standard conditions and exposed for 24 h to an “active” or to a mock BR device, in resting conditions and during treatment with H_2_O_2_ at the concentration of 500 μM, added at the beginning of the 24 h period. The personnel who performed the experiments, collected, and analysed the data was unaware of which device was “active” and which was mock. At the end of the culture, HUVEC were harvested and evaluated for viability and mitochondrial metabolic activity by means of the Trypan Blue exclusion and the 3-(4,5-dimethyl-2-thiazolyl)-2,5-diphenyl-2H tetrazolium bromide reduction method, respectively.

Under control conditions, viability and mitochondrial metabolic activity were not different in HUVEC exposed to the “active” or to the mock device. HUVEC viability, however, was significantly reduced by exposure to H_2_O_2_ in samples exposed to the mock device but not in those exposed to the “active” device. HUVEC mitochondrial metabolic activity was significantly reduced by exposure to H_2_O_2_ in both samples exposed to the “active” and to the mock device, however reduction was significantly less in samples exposed to the “active” device.

In conclusion, exposure to the BR-based device QDOME MINI PERSONAL, produced by the society QMED SWISS SA (Lugano, CH), prevented the H_2_O_2_-induced reduction of viability and reduced the H_2_O_2_-induced impairment of mitochondrial metabolic activity in cultured HUVEC.

## INTRODUCTION

Bioresonance (BR) can be defined as ‘a method … which employs the ‘patients’ own vibrations’; the aim is to measure biophysical vibrations (frequencies) of a patient … and subsequently feed them back into the body in modified form via a second electrode’. Essentially BR claims that, with sophisticated electronic devices (of which several modifications exist), electromagnetic waves can be measured (the diagnostic element) and, if abnormal, be normalised (the therapeutic element of the approach). BR approaches have been heavily criticized because of their use of pseudoscientific language and the lack of evidence supporting their effectiveness (Ernst, 2004), however the medical and scientific literature includes several studies claiming benefits from BR techniques in a heterogeneous number of diseases, such as: depression (Muresan et alo., 2021), cigarette smoking (Pihtili et al., 2014), diabetes mellitus (Jalan et al., 2022), overtrained athlete syndrome (Badtieva et al., 2018), gonarthrosis (Maĭko and Gogoleva, 2000), irritable bowel syndrome-associated low-back pain (Barassi et al., 2024), lymphedema and lipedema (Elio et al., 2014; Cavezzi et al., 2013), “non-organic” gastrointestinal complaints (Nienhaus and Galle, 2006), chronic psoriasis vulgaris (Del Giudice et al., 2015), and even for improvement of well-being in healthy individuals (Walach and Marmann, 2023). Unfortunately, most of these are low-quality studies and results are often not reproducible when subject to rigorous testing, for example in atopic dermatitis in children (Schöni et al., 1997), in the diagnosis of allergy (Wüthrich, 2005).

Nonetheless, circumstantial evidence also exists showing effects in animal models of diseases, for example tumor regression in rats inoculated with Morris hepatoma cells (Fedorowski et al., 2004), and direct functional effects in human cells and tissues, such as: increased protein synthesis in lymphocytes (Islamov et al., 1995), complex biological effects in lung and liver cells (Podchernyaeva et al., 2008), increased intracellular antioxidant systems in lymphocytes from rheumatoid arthritis patients (Islamov et al., 2002), stimulation of metabolism in cultured fibroblasts (Fraunhofer Institute, 2011), stimulation of healing in cultured fibroblasts and regeneration of intestinal epithelial cells (Dartsch Scientific GMBH).

The present study was devised to assess the possible effects of BR-based devices produced by the society QMED SWISS SA (Lugano, CH) on the viability and metabolic activity of human umbilical venous endothelial cells (HUVEC).

## MATERIALS AND METHODS

### BR-based devices

The following BR device was chosen for the experiments: QDOME MINI PERSONAL - measures: 5 X 5 CM (Figure 1). QMED SWISS SA provided an “active” device and a mock device, each marked with a different letter (A and B). The personnel who performed the experiments, collected and analysed the data was unaware of which device was “active” and which was mock. QMED SWISS SA revealed which was the “active” device only after collection and analysis of the data were concluded and the results were shared and discussed.

**Figure 1.**
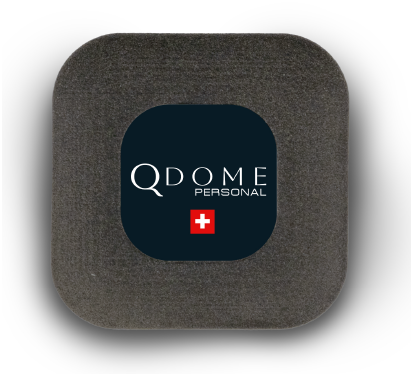
QDOME MINI PERSONAL (https://qdome.ch/en/prodotti/harmonizers/qdome-personal-mini/).

The devices were handled and used according to the written indications provided by QMED SWISS SA. QMED SWISS SA also provided a cover of its special fabric to shield the incubator in which the experiments took place (Figure 2).

**Figure 2.**
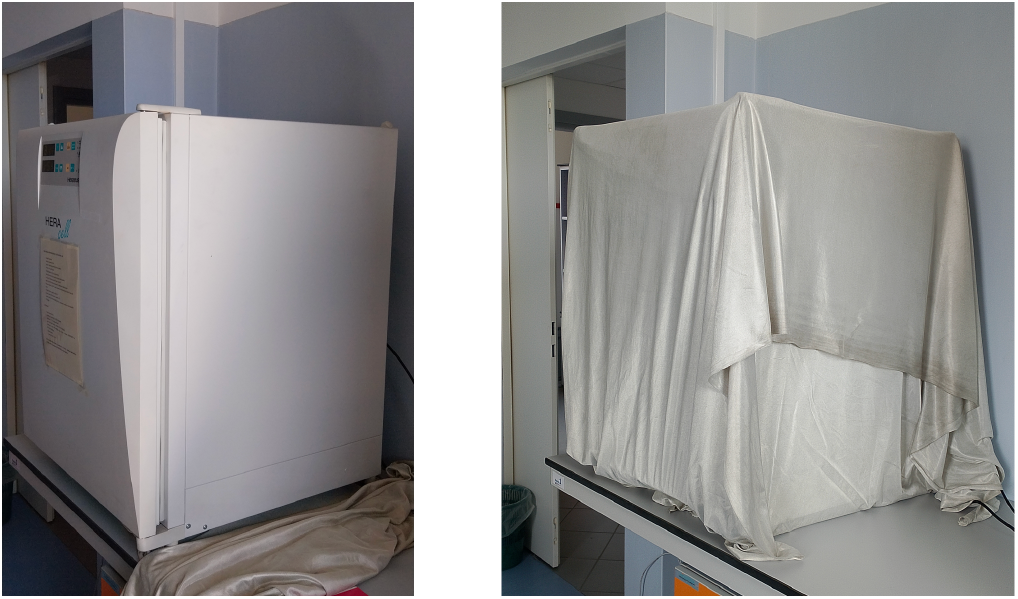
Cell culture incubator in which the experiments took place, without (left) and with the special cover provided by QMED SWISS SA (right).

### HUVEC culture

Experiments were performed on human umbilical vein endothelial cells (HUVEC). HUVEC were purchased from PromoCell (cod. C-12205, PromoCell GmbH) and routinely cultured in endothelial cell growth medium kit (cod. C-22110, PromoCell GmbH) containing fetal calf serum (FCS) (0,02 ml/ml), endothelial cell growth supplement/heparin (ECGS/H) (0,004 ml/ml), human epithelial growth factor (hEGF) (0,1 ng/ml), human basic fibroblast growth factor (hb-FGF) (1 ng/ml), and hydrocortisone (1 µg/ml) (Endothelial Cell Growth Complete Medium). The culture flasks were maintained at 37°C in a humidified atmosphere of 5% CO_2_. HUVEC used for the experiments were between passage 11 and 15.

### Treatments

Confluent cells were exposed for 24 h to the “active” or to the mock BR device in resting conditions and during treatment with H_2_O_2_ (29-32% w/w hydrogen peroxide in water, cod. 216763, Sigma-Aldrich, Saint Louis, Missouri, USA) at the concentration of 500 μM, added at the beginning of the 24 h period. At the end of the culture, HUVEC were evaluated for viability and mitochondrial metabolic activity.

### Viability

The viability of HUVEC was assessed by the Trypan Blue exclusion test using a Cellometer Auto T4 Cell Counter (Nexcelom Bioscience LLC, Euroclone). To this end, HUVEC cells were resuspended in Endothelial Cell Growth Complete Medium. Cells were then seeded in a 48-well flat bottom plate at a density of 1,5 x 10^4^ cells/well (final volume: 500 µl) and cultured for 24 h at 37 °C in a 5% CO_2_ atmosphere. After this period, HUVEC formed a monolayer that was left untreated or treated with H_2_O_2_ 500 µM for another 24 h. Finally, cells were detached using trypsin-EDTA 1X in phosphate buffered saline (PBS) (cod. ECB3052D, Euroclone), centrifuged and pellets were resuspended in 100 µl of medium. Aliquots of 20 µl of each sample were stained with 20 µl of trypan blue solution and 20 µl of this cell suspension were loaded into the Cellometer counting chamber. The viability was measured as both % of viable cells and as total cell count.

### Mitochondrial metabolic activity

Mitochondrial metabolic activity in HUVEC cells was assessed by means of the MTT [3-(4,5-dimethyl-2-thiazolyl)-2,5-diphenyl-2H tetrazolium bromide] reduction method (Mosmann, 1983). In short, HUVEC cells were resuspended in Endothelial Cell Growth Complete Medium. Cells were then seeded in a 96-well flat bottom plate and cultured for 24 h at 37 °C in 5% CO_2_ atmosphere. The absorbance (optical density, OD, in arbitrary units) was measured using a microplate spectrophotometer (Thermo Fisher Scientific Multiskan FC, Thermo Fisher Scientific Inc.) with a 570 nm test wavelength and a 650 nm reference wavelength. Results were expressed as mean OD value of duplicates after subtracting the absorbance of the culture medium alone.

### Statistics

Data are shown as means±standard deviation (SD) of the mean, unless otherwise indicated, with n showing the number of replicates. Unless otherwise specified, two-way ANOVA was used to test the statistical significance of differences between groups. GraphPad Prism 10 for macOS Version 10.5.0 (673), May 27, 2025 (GraphPad Software, La Jolla California USA, www.graphpad.com) was used to perform statistical analyses and for the preparation of graphic figures.

## RESULTS

### Viability

The Trypan Blue exclusion test was performed on 32 different HUVEC preparations, 16 exposed to device A and 16 to device B. Each preparation was used to prepare two samples, one kept under control conditions, and one treated with H_2_O_2_ 500 µM. Finally, cell viability was successfully measured in all but 6 samples, 4 exposed to device A and 2 to device B, where assessment of percentage viable cells and cell count could not be performed due to technical/analytical reasons. HUVEC used for the experiments were between passage 12 and 15 (median: 13,5) with device A, and between passage 12 and 15 (median: 14) with device B (P = 0,462 by two-tailed Student’s t test for unpaired values, n = 16).

Under control conditions, the percentages of viable cells and cell counts were not different between samples exposed to device A and those exposed to device B (Table 1 and Figure 3). In samples treated with H_2_O_2_ 500 µM, however, the percentage of viable cells and cell count were significantly reduced only in samples exposed to device A, while neither viable cells nor cell count were different from control conditions in samples exposed to device B (Table 1 and Figure 3).

**Table 1.** Effect of BR devices on HUVEC viability assessed by the Trypan Blue exclusion test. The viability is expressed as % of viable cells and as total cell count. Data are shown as means±SD of n = 14-15 replicates.

**1. Viability (%)**
| | alone | + H<sub>2</sub>O<sub>2</sub> 500 μM | P<br>vs respective cells alone |
| --- | --- | --- | --- |
| **Device A** | 92,5±4,5 | 84,3±9,8 | **0,0004** |
| **Device B** | 91,9±3,2 | 91,0±4,1 | 0,6573 |
| **P**<br>Device A vs Device B | 0,8005 | **0,0029** |  |

**2. Cell count (n x 10.000)**
| | alone | + H<sub>2</sub>O<sub>2</sub> 500 μM | P<br>vs respective cells alone |
| --- | --- | --- | --- |
| **Device A** | 1,90±0,55 | 1,39±0,66 | **0,0124** |
| **Device B** | 1,78±0,43 | 1,80±0,51 | 0,9263 |
| **P**<br>Device A vs Device B | 0,5327 | **0,0422** |  |

**Figure 3.**
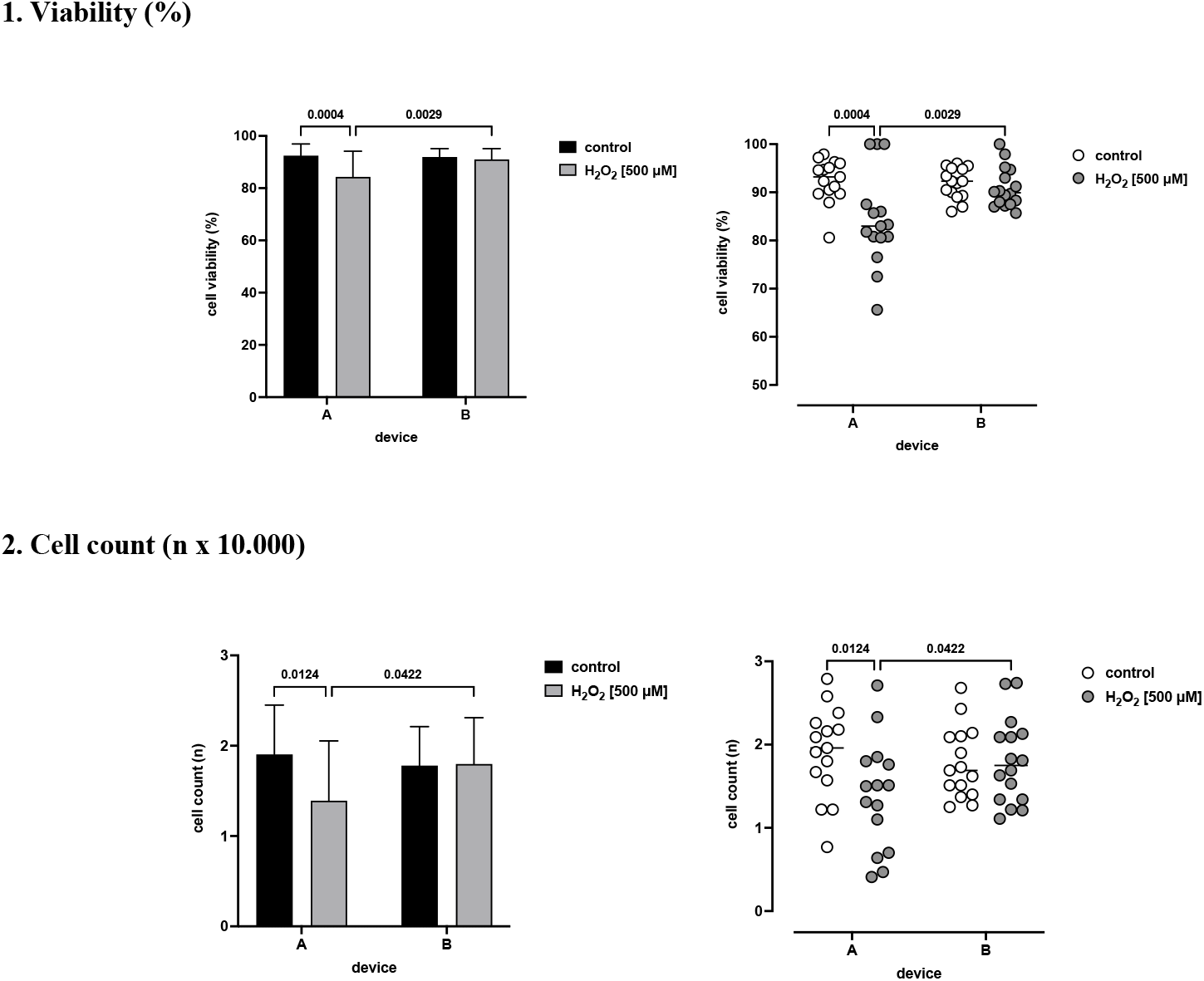
Effect of BR devices on HUVEC viability assessed by the Trypan Blue exclusion test. The viability is expressed as % of viable cells and as total cell count. Data are shown as means±SD of n = 14-15 replicates (left panels) and as values of individual experiments (right panels).

### Mitochondrial metabolic activity

The MTT assay was performed on 96 different HUVEC preparations, 48 exposed to device A and 48 to device B. As with the Trypan Blue exclusion test, each preparation was split in two samples, one kept under control conditions, and one treated with H_2_O_2_ 500 µM. Finally, mitochondrial metabolic activity was successfully measured in all the samples. HUVEC used for the experiments were between passage 11 and 15 (median: 12,5) with device A, and between passage 11 and 15 (median: 13) with device B (P = 0,080 by two-tailed Student’s t test for unpaired values, n = 48).

Under control conditions, mitochondrial metabolic activity was not different between samples exposed to device A and those exposed to device B (Table 2 and Figure 4). Treatment with H_2_O_2_ 500 µM significantly reduced mitochondrial metabolic activity with both device A and device B, however to different extents. In samples exposed to device A, metabolic activity was reduced by about 63%, while in samples exposed to device B metabolic activity was reduced only by 43%, a difference which was highly significant (P<0,0001) (Table 2 and Figure 4).

**Table 2.**
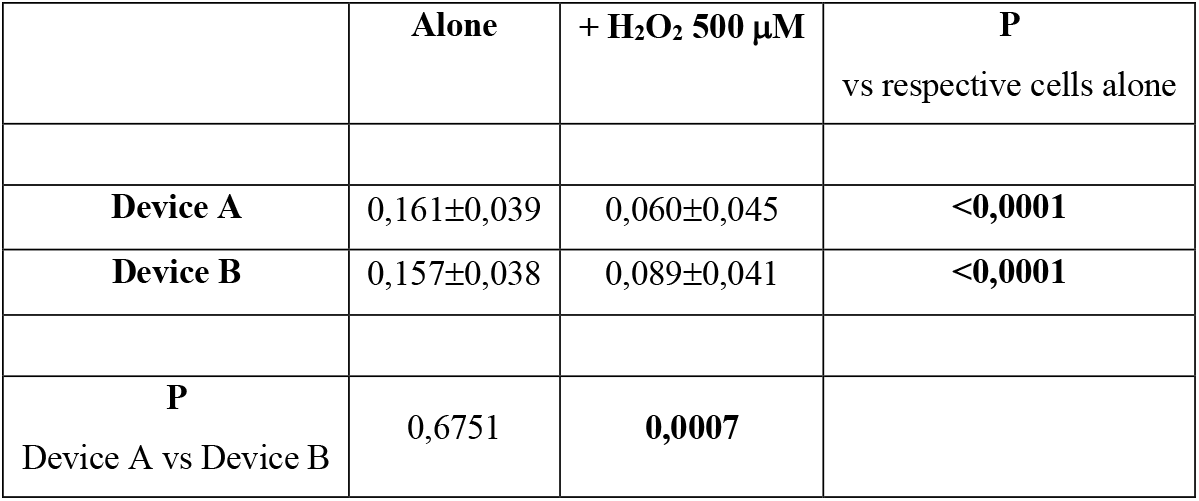
Effect of BR devices on HUVEC mitochondrial metabolic activity assessed by the MTT reduction method. The mitochondrial metabolic activity is expressed as mean OD value (in arbitrary units) of duplicates after subtracting the absorbance of the culture medium alone. Data are shown as means±SD of n = 48 replicates.

**Figure 4.**
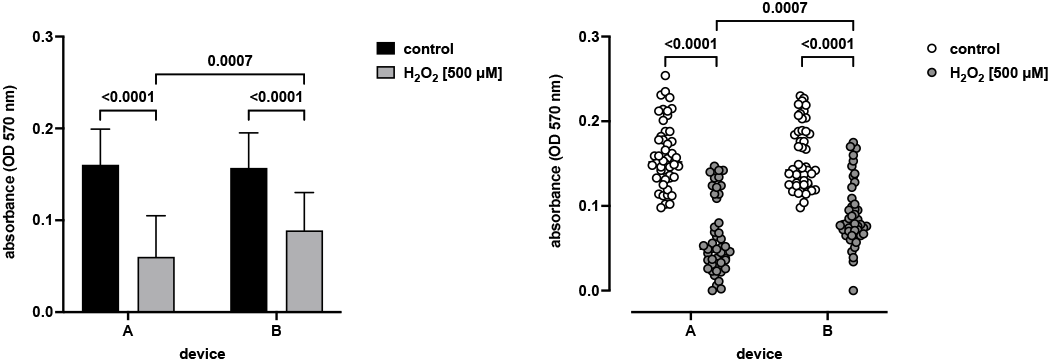
Effect of BR devices on HUVEC mitochondrial metabolic activity assessed by the MTT reduction method. The mitochondrial metabolic activity is expressed as mean OD value (in arbitrary units) of duplicates after subtracting the absorbance of the culture medium alone. Data are shown as means±SD of n = 48 replicates (left panel) and as values of individual experiments (right panel).

## CONCLUSIONS

The present study has been undertaken to assess whether the BR-based device QDOME MINI PERSONAL, produced by the society QMED SWISS SA (Lugano, CH), could affect the viability and mitochondrial metabolic activity of HUVEC. To this end, QMED SWISS SA provided a “active” device and a mock device, each marked with a different letter (A and B). The study was brought about by personnel who was unaware of which device was “active” and which was mock. QMED SWISS SA revealed which was the “active” device only after collection and analysis of the data were concluded and the results were shared and discussed.

The results of the study can be summarised as follows:

1. under control conditions, viability and mitochondrial metabolic activity were not different in HUVEC exposed to device A or to device B;
2. HUVEC viability was significantly reduced by exposure to H_2_O_2_ only in samples exposed to device A but not in those exposed to device B;
3. HUVEC mitochondrial metabolic activity was significantly reduced by exposure to H_2_O_2_ in both samples exposed to device A and to device B, however reduction was significantly less in samples exposed to device B.

While the effect on mitochondrial metabolic activity could imply either that device B is protective or that device A is detrimental to HUVEC, the observation that HUVEC viability is reduced by H_2_O_2_ only in the presence of device A but not of device B strongly suggests that the best overall explanation for the whole set of results is that device B protects HUVEC from H_2_O_2_-induced damage. This interpretation is also supported to some extent by the observation that HUVEC viability and mitochondrial metabolic activity were not affected under control conditions. Most importantly, this interpretation of the results is consistent with the fact that the “active” device is device B, supporting the conclusion that the effects on HUVEC viability and metabolic activity are related to the activity of the device.

In this study, HUVEC were examined by the Trypan Blue exclusion test and by the MTT assay. The Trypan Blue exclusion test is a classical reference method developed in 1975 to determine cell viability (Avelar-Freitas et al., 2014; Tran et al., 2011; Kamiloglu et al., 2020). Trypan blue is an azo dye, derived from toluidine, which does not cross intact cell membranes. Consequently, Trypan Blue is excluded from viable cells (Abhishek et al., 2018). On the contrary, Trypan Blue dye penetrates cell membrane of necrotic cells due to loss of cell membrane integrity and enters the cytoplasm. Thus, under light microscope, only necrotic (dead) cells absorb the dye and are stained in blue colour (Chan et al., 2020). On the other side, the MTT reagent, a mono-tetrazolium salt consisting of a positively charged quaternary tetrazole ring core containing four nitrogen atoms surrounded by three aromatic rings including two phenyl moieties and one thiazolyl ring, can be reduced resulting in disruption of the core tetrazole ring and formation its violet-blue water-insoluble formazan (Berridge et al., 2005). MTT easily crosses the cell membrane and the mitochondrial inner membrane of viable cells, and is reduced to its formazan by metabolically active cells (Berridge et al., 2005; Stockert et al., 2018). Mosmann et al. in 1983 exploited the chromogenic nature of this redox chemical reaction to develop a colorimetric-based assay of cell metabolic activity (Mosmann, 1983).

On this basis, it is possible to argue that device B protects HUVEC from the oxidative damage induced by exposure to H_2_O_2_ both in terms of maintaining membrane integrity and supporting cellular metabolism, particularly mitochondrial metabolism. On the other side, the present study provides no information about the mechanisms responsible for such effects or about the molecular pathways involved. In this regard, further experiments are warranted to investigate for example the intracellular pathways involved in the response to oxidative stress and in the regulation of cell viability and apoptosis.

## Acknowledgements

This study was supported by a grant from QMED SWISS SA (Lugano, CH) - https://qmedswiss.ch/

